# Insulin-like Growth Factor 2 mRNA-binding protein 2 (IGF2BP2) Promotes Castration-Resistant Prostate Cancer Progression by Regulating AR-V7 mRNA Stability

**DOI:** 10.1101/2024.04.07.588211

**Authors:** Taruna Saini, Devesh Srivastava, Rajnikant Raut, Parul Mishra, Ashish Misra

## Abstract

The emergence of constitutively active androgen receptor (AR) splice variant AR-V7 poses a formidable challenge in treating prostate cancer, as it lacks the ligand binding region targeted by androgen deprivation therapies such as enzalutamide and abiraterone. AR-V7 is critical for castration-resistant prostate cancer (CRPC) development and progression, however the molecular mechanisms regulating its expression and biological function remain poorly understood. Here, we investigate the role of IGF2BP2 in regulating AR-V7 expression and CRPC progression. We demonstrate that IGF2BP2 silencing leads to downregulation of AR-V7 and its downstream target genes without affecting AR levels. Additionally, IGF2BP2 knockdown also enhances the sensitivity of CRPC cells to enzalutamide while overexpression increases AR-V7 expression and confers increased resistance to enzalutamide. Mechanistically, our experiments demonstrate that IGF2BP2 binds to the intronic splicing enhancer (ISE) region of AR-V7, thereby enhancing its mRNA stability Furthermore, our domain-deletion analysis pinpoints the role of KH3 and KH4 domains of IGF2BP2 in regulating AR-V7 stability and enzalutamide resistance. Taken together, our findings suggest that IGF2BP2 plays a critical role in regulating AR-V7 expression and stability, offering a novel target for developing therapeutic interventions for CRPC.

## 1. Introduction

Prostate cancer is the second deadliest cancer among men worldwide. The growth and survival of prostate cancer depends on the steroid nuclear receptor termed androgen receptor (AR). It belongs to the hormone receptor family, and comprises of 8 exons that encode for N-terminal, DNA binding, hinge region, and the ligand binding domains (1). Androgen deprivation therapy (ADT), employing medications such as abiraterone and enzalutamide (2), serves as an AR-targeted approach to impede the advancement of prostate cancer. Abiraterone impairs AR signaling by modulating androgen synthesis, while enzalutamide blocks AR activity by binding to LBD. Although initially effective, patients quickly develop resistance to these therapies, leading to the emergence of CRPC (3). While multiple pathways are known to contribute to CRPC progression, the emergence of androgen receptor splice variant, AR-V7, is believed to be the key driver of CRPC (4–7). Clinical studies have unequivocally demonstrated that increased AR-V7 expression relates to poor prognosis in CRPC patients (8,9).

Numerous post-transcriptional regulators have emerged as pivotal players in regulating AR-V7 generation and activity in CRPC, offering promising avenues for therapeutic intervention. Among these regulators, RNA-binding proteins (RBPs) are known to play a critical role in modulating mRNA processing, polyadenylation, stability, and translation (10,11). Extensive studies have underscored the importance of RBPs in CRPC development and progression (12,13). The expression of heterogeneous nuclear ribonucleoprotein A1 (hnRNPA1) and Sam68 positively correlates with AR-V7 generation and expression in prostate cancer (14,15). In fact, the molecular circuitry involving hnRNPA1, NF-κB2/p52 and c-Myc, intricately modulates AR-V7 generation in prostate cancer cells (14). hnRNPH1 and hnRNPL have also been shown to affect AR splicing and contribute to CRPC development (16,17). PTB-associated splicing factor (PSF) has been shown to promote generation of AR and its splice variants by modulating the spliceosome machinery (18). Therefore, the discovery of previously undocumented RBPs driving AR-V7 expression will provide alternative treatment options for CRPC.

IGF2BP2 is a member of the IGF2 mRNA-binding protein family, that regulate various post-transcriptional processes such as mRNA localisation, stability, and translation. IGF2BP2 is a 66kDa protein comprising of two-RRMs domains at the N-terminal and four-hnRNP-K homology (KH) domains at the C-terminus (19). Studies have shown that IGF2BP2 dysregulation is associated with a variety of diverse diseases including cancer, diabetes and insulin resistance (19). Overexpression of IGF2BP2 is known to promote tumour progression in a variety of malignancies such as glioblastoma and colorectal cancer (20–22) . Mounting evidence also suggests that IGF2BP2 acts as a m6A reader, recognizing the N6-methyladenosine (m6A) modifications on transcripts critical for cancer initiation and progression (23,24).

Here, we demonstrate that IGF2BP2 is significantly upregulated in CRPC patients and promotes CRPC cell proliferation by regulating AR-V7 expression. Knockdown of IGF2BP2 results in downregulation of AR-V7 and its downstream target genes. Furthermore, our experiments highlight the involvement of IGF2BP2 in proliferation of CRPC cells, stemness properties as well as tumorigenesis and establish a link between IGF2BP2 expression and enzalutamide resistance in CRPC cells. Mechanistically, we show that IGF2BP2 binds directly to ISE region of AR-V7 pre-mRNA via its KH3 and KH4 domains and plays a critical role in maintaining AR-V7 mRNA stability.

## 2. Results

### 2.1. IGF2BP2 regulates AR-V7 expression

To determine the clinical relevance of IGF2BP2 in CRPC, we analysed the mRNA expression data for prostate cancer patients available in the TCGA database, encompassing normal prostate (N=52), primary prostate cancer (N=498), and CRPC (N=99) samples. Our analysis revealed that IGF2BP2 is significantly upregulated in CRPC patient samples compared to primary prostate cancer samples (*p* <L0.01, logFC= 1.88) (Fig. 1A). We also observed that IGF2BP2 is upregulated in CRPC patient samples relative to the normal prostate samples (logFC=0.7, p value=0.031) (Fig. S1). These results point towards positive correlation between IGF2BP2 expression in patient samples and CRPC progression.

**Figure 1:**
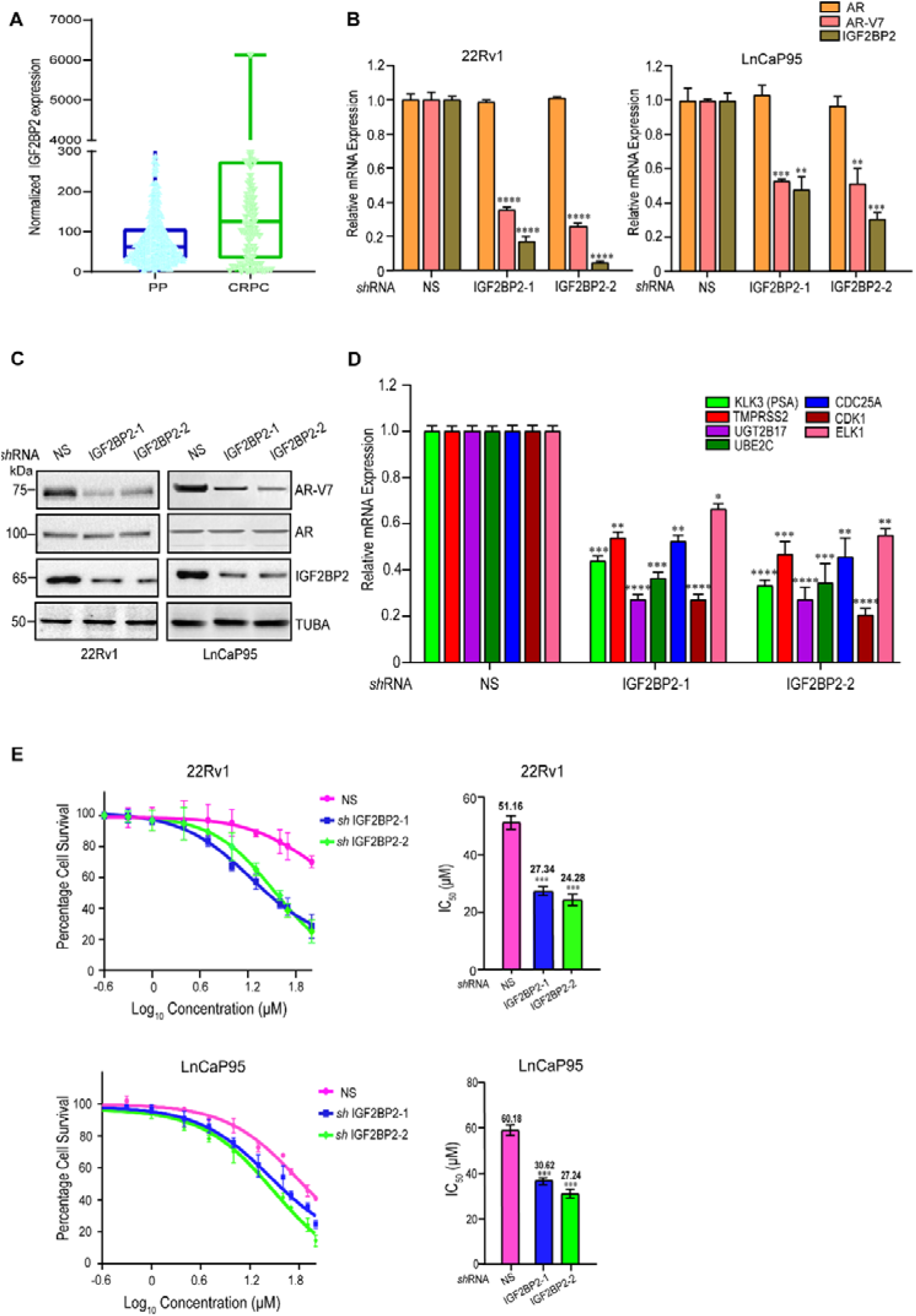
IGF2BP2 regulates AR-V7 expression. **A.** IGF2BP2 mRNA expression in primary prostate (PP) (n= 500) compared to CRPC (n=99) patient samples from TCGA dataset (*p* < 0.01, logFC= 1.88). mRNA expression is graphically represented as box and whiskers plot with outliers. **B.** qRT-PCR analysis of AR-V7, AR and IGF2BP2 mRNA expression in 22Rv1 and LnCaP95 cell lines expressing NS or IGF2BP2 shRNAs. **C.** Immunoblot analysis of AR-V7 and AR in 22Rv1 and LnCaP95 cell lines expressing NS or IGF2BP2 shRNAs. Tubulin was used as the loading control. **D.** qRT-PCR analysis of indicated genes upon NS or IGF2BP2 knockdown in 22Rv1 cells. **E.** MTT assay to determine IC50 of enzalutamide in 22Rv1 and LnCaP95 cells upon IGF2BP2 or NS knockdown. The data are shown as the mean ± SD; *n* = 3 independent experiments, two-tailed Student’s t-test, \**p* < 0.05, \*\**p* < 0.01, \*\*\**p* < 0.001.

Next, to probe the role of IGF2BP2 in AR-V7 expression, we analysed the expression of AR-V7 and AR in LnCaP95 and 22Rv1 cells infected with two sequence-independent shRNAs against IGF2BP2 (Fig. 1B). Cells expressing non-silencing (NS) were used as control. We observed that downregulation of IGF2BP2 resulted in a significant decrease in AR-V7 expression in both qRT-PCR (Fig. 1B) and immunoblot analysis (Fig. 1B). Notably, full-length AR levels remained unchanged, pointing towards the specificity of IGF2BP2 in regulating AR-V7 levels (Fig. 1A and B).

AR-V7 is a constitutively active AR-variant that regulates the canonical AR target genes, along with an unique set of genes such as UGT2B17, UBE2C, CDC25A, CDK1, and ELK1 (25–27). We examined the impact of IGF2BP2 knockdown on these target genes using real-time PCR (qRT-PCR). Our results demonstrate that the expression of AR-V7-regulated genesUGT2B17, UBE2C, CDC25A, CDK1, ELK1, and prostate-specific antigen (PSA)/KLK3, is significantly decreased upon IGF2BP2 knockdown (Fig. 1D) suggesting that alterations in IGF2BP2 expression directly affect the expression of AR-V7 target genes.

Next, to examine the impact of IGF2BP2 on enzalutamide resistance, we measured the IC50 of enzalutamide in IGF2BP2 knockdown 22Rv1 and LnCaP95 cells compared to non-silencing (NS) shRNA infected cells using MTT assay. Knockdown of IGF2BP2 in 22Rv1 and LnCaP95 cells led to substantial decrease in IC50 of enzalutamide relative to NS knockdown cells. The IC50 for enzalutamide decreased from 51.16μM in NS-infected 22Rv1 cells to 27.34μM and 24.28μM in IGF2BP2 knockdown 22Rv1cells. In LnCaP95 cells, the IC50 decreased from 60.18μM NS-infected cells to 30.62μM and 27.24μM in IGF2BP2 knockdown cells (Fig. 1E). Collectively, our results demonstrate the role of IGF2BP2 in regulating AR-V7 expression and enzalutamide resistance in CRPC cells.

### 2.2 IGF2BP2 recruitment to specific splice sites on AR pre-mRNA is required to maintain its stability

Under androgen-deprivation conditions, distinct splice sites on AR mRNA associate with various splicing factors and RNA-binding proteins to regulate alternative splicing. Notably, ISE (intronic splicing enhancer) and ESE (exonic splicing enhancer) sites ( near the 3′ splice site of exon 3b play a pivotal in AR-V7 generation (28,29) (Fig. 2A). To determine, if IGF2BP2 is recruited to these splice sites, we performed an RNA immunoprecipitation assay followed by qRT-PCR (RIP-qPCR) using anti-FLAG antibody in both FLAG-IGF2BP2 and vector-expressing VCaP cells. The RIP-qPCR assay results revealed that IGF2BP2 is highly enriched in the ISE region compared to ESE region of AR-V7 pre-mRNA (Fig. 2B and Fig. S2).

**Figure 2:**
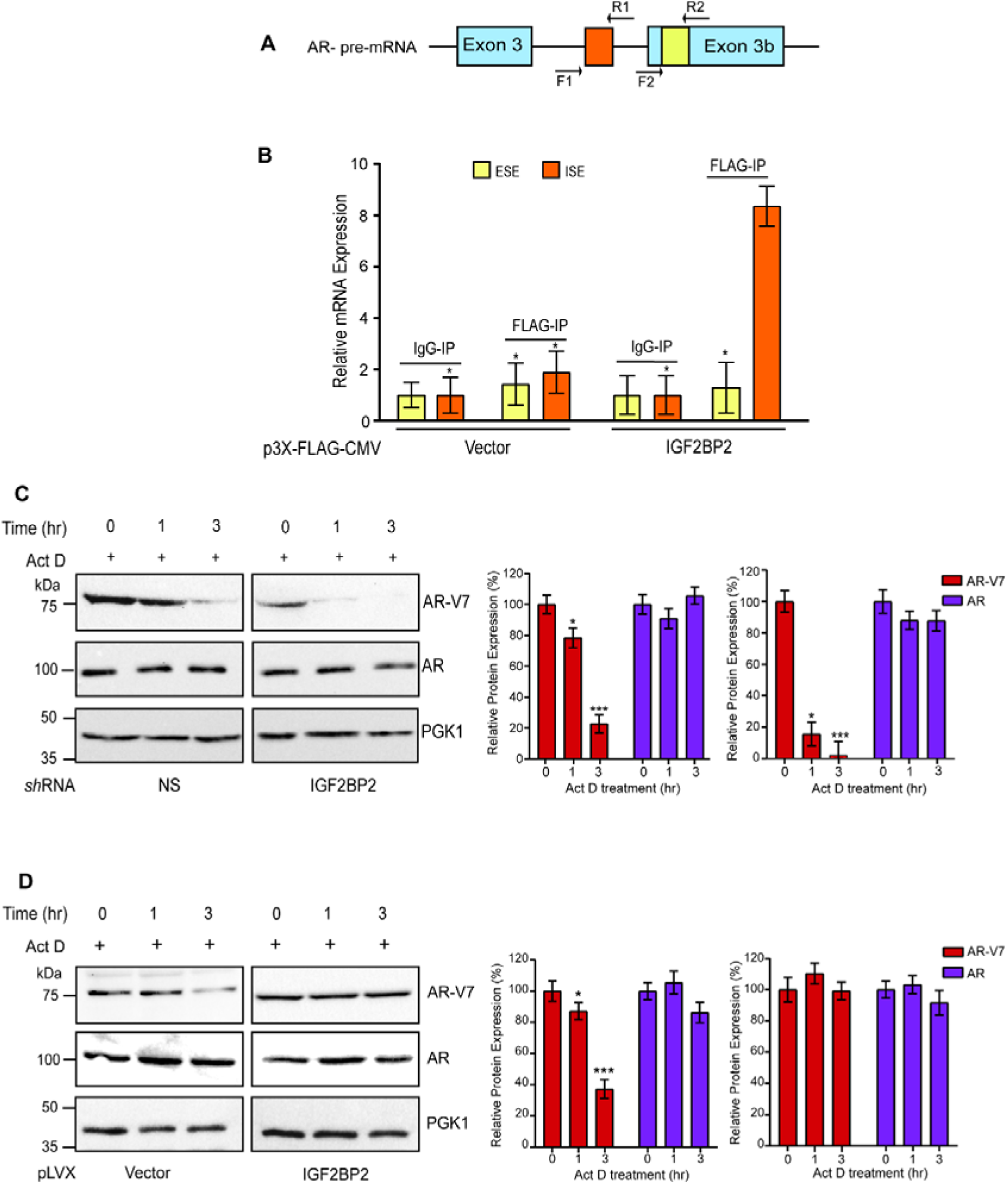
Recruitment of IGF2BP2 to specific splice sites on AR pre-mRNA is required to maintain stability. **A.** Schematic representation of ISE and ESE sites on exon 3b of AR pre-mRNA. F1-R1 and F2-R2 represent the qPCR primer locations in the ISE and ESE regions. **B.** RIP-qPCR analysis showing enrichment of IGF2BP2 on ISE and ESE regions of AR-V7 mRNA in IGF2BP2-overexpressing VCaP cells relative to vector-overexpressing cells. IgG was used as negative control. Fold-enrichment relative to IgG for indicated sites is shown **C.** Immunoblot analysis of indicated proteins in 22Rv1 cells expressing IGF2BP2 or NS shRNAs following actinomycin-D (5μg/mL) treatment for 0, 1 and 3hr. (n=3 biological replicates/group) **D.** Immunoblot analysis of indicated proteins in IGF2BP2-overexpressing VCaP cells relative to vector-overexpressing cells following actinomycin-D (5μg/mL) treatment for 0, 1 and 3hr. (n=3 biological replicates/group). Densitometric analysis for relative AR-V7 and AR expression was performed using Image J software and normalised to PGK1 expression. Data are the mean± SD. \**p* < 0.05, \*\**p* < 0.01, *** *p* < 0.001.

Next, to understand the effect of IGF2BP2 binding to AR-V7 mRNA, we investigated the role of IGF2BP2 in regulating the stability of AR-V7. We assessed the half-life of AR-V7 protein upon actinomycin D treatment in 22Rv1 cells expressing either NS or IGF2BP2 shRNAs. Our immunoblot results revealed that AR-V7 levels are significantly reduced in cells expressing IGF2BP2 shRNA relative to NS expressing cells 3 hr post actinomycin D treatment, while no effect is observed on AR levels (Fig. 2C, Fig. S3). This is consistent with enhanced AR-V7 stability in VCaP cells expressing IGF2BP2 relative to vector-expressing cells after actinomycin D treatment (Fig. 2D). Collectively, our results demonstrate that IGF2BP2 directly binds to the ISE region of AR-V7 mRNA and plays a crucial role in enhancing its stability.

### 2.3 IGF2BP2 promotes migration and stemness in CRPC cells

Cancer progression is marked by the ability of malignant cells to migrate and invade surrounding tissues (30). We performed wound-healing assay in NS and IGF2BP2 shRNA expressing 22Rv1 cells to assess the potential of IGF2BP2 in regulating migration properties of CRPC cells. Our results revealed a ∼40-50% reduction in cell migration upon IGF2BP2 knockdown compared to NS (Fig. 3A). Next, we assessed the effect of IGF2BP2 on colony formation ability of CRPC cells. Our results demonstrate that there is a substantial decrease in the number of colonies in 22Rv1 and LnCaP95 cell lines expressing IGF2BP2-shRNAs compared to NS (Fig. 3b). Moreover, reduced expression of IGF2BP2 is accompanied by the loss of expression of stemness- and EMT-related genes such as Sox-2, Oct-4, NANOG, E-cadherin and Vimentin (Fig. S4). These findings suggest a positive correlation between IGF2BP2 expression, stemness and the-colony-formation ability of CRPC cells. Next, we evaluated the ability of IGF2BP2 knockdown CRPC cell lines 22Rv1 and LnCaP to grow in an anchorage independent manner using soft agar colony formation assays. NS-shRNA expressing CRPC cells were used as negative control. Our results demonstrate that IGF2BP2 knockdown leads to significant decrease in the ability of CRPC cells to form colonies in soft agar (Fig. 3c).

**Figure 3:**
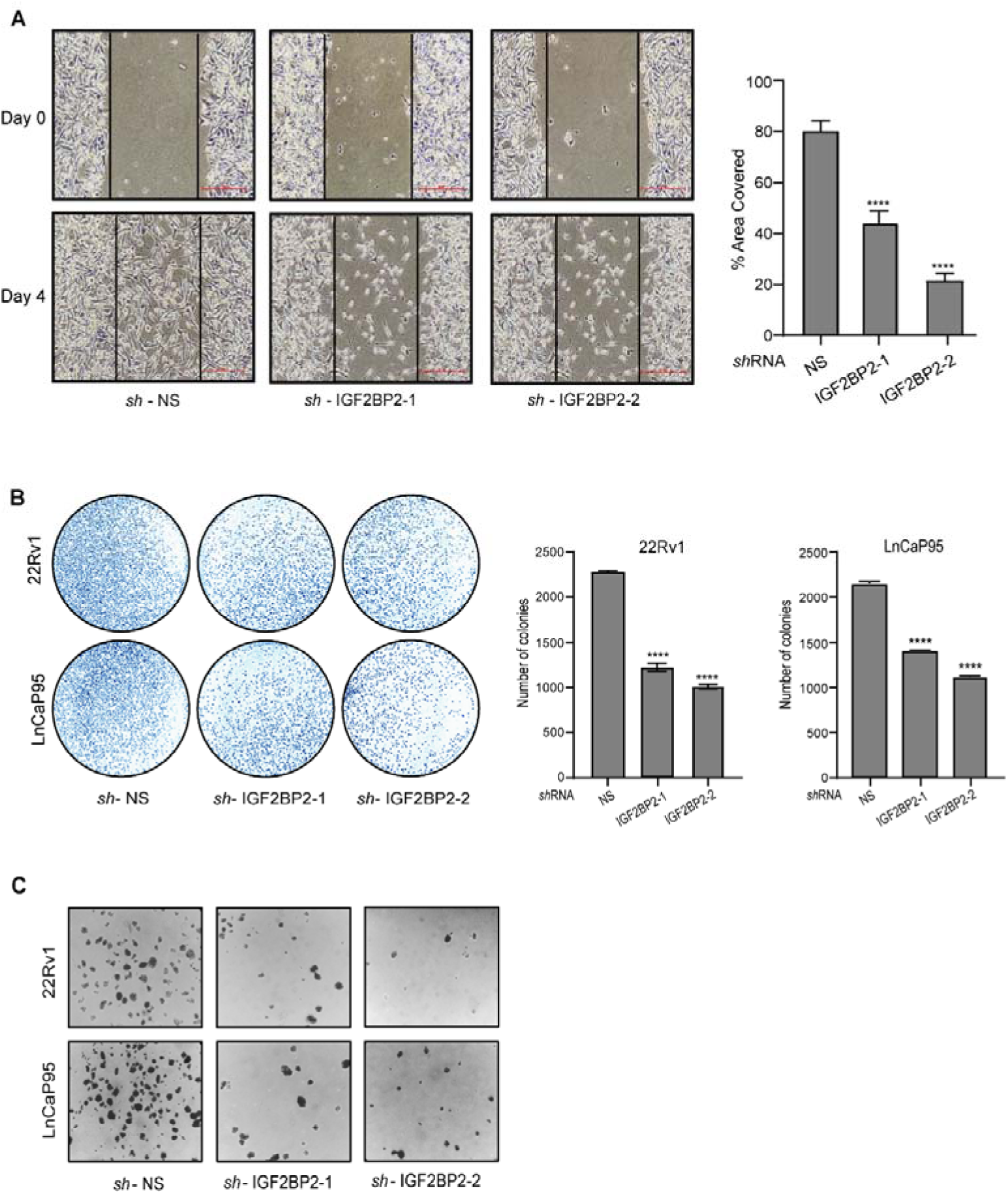
IGF2BP2 promotes migration and metastasis in CRPC cells. **A.** Wound healing assay in 22Rv1 cells expressing IGF2BP2 or NS shRNAs. Quantification of the total area covered by the cells was calculated in ImageJ software and plotted (**** *p* < 0.0001). Scale bar, 100μm. **B.** Colony formation assay in 22Rv1 and LnCaP95 cells expressing IGF2BP2 or NS shRNAs. Colonies were stained using crystal violet and counted (n=3 biological replicates/group). Error bars indicate standard deviation and **** *p* < 0.0001 **C.** Representative stained images showing soft agar colony formation in 22Rv1 and LnCaP95 cell lines expressing IGF2BP2 or NS shRNAs. Images were taken after 21 days.

Collectively, the above results highlight the involvement of IGF2BP2 in regulating CRPC cell migration, proliferation, colony formation, and anchorage-independent growth, underscoring its role as a key player in CRPC progression.

### 2.4 IGF2BP2 overexpression enhances AR-V7 expression and enzalutamide resistance

Next, we investigated the impact of IGF2BP2 overexpression on tumorigenic properties of VCaP cells. We infected VCaP cells with constructs overexpressing either empty vector or FLAG-IGF2BP2 and monitored the levels of both AR and AR-V7 using both qRT-PCR and immunoblotting. Our results demonstrated that IGF2BP2 overexpression specifically enhances AR-V7 levels inside VCaP cells while no effect was observed on full-length AR levels (Fig. 4A and B). Additionally, the expression of AR-V7 downstream genes, including UGT2B17, UBE2C, CDC25A, CDK1, and ELK1, was found to be markedly elevated in IGF2BP2-overexpressing cells compared to vector-expressing cells (Figure 4b). Consistent with these molecular changes, we also observe increase in colony forming ability of cells overexpressing IGF2BP2 relative to vector-expressing cells (Fig. 4C). Next, we assesed the impact of IGF2BP2 overexpression on enzalutamide sensitivity of VCaP cells using MTT assay. Our results demonstrate that IGF2BP2-overexpression enhances enzalutamide IC50 value from 19.4μM in vector-expressing cells to 44.5μM in IGF2BP2-overexpressing cells, suggesting the potential contribution of IGF2BP2 in conferring enzalutamide resistance to CRPC cells (Fig. 4D).

**Figure 4:**
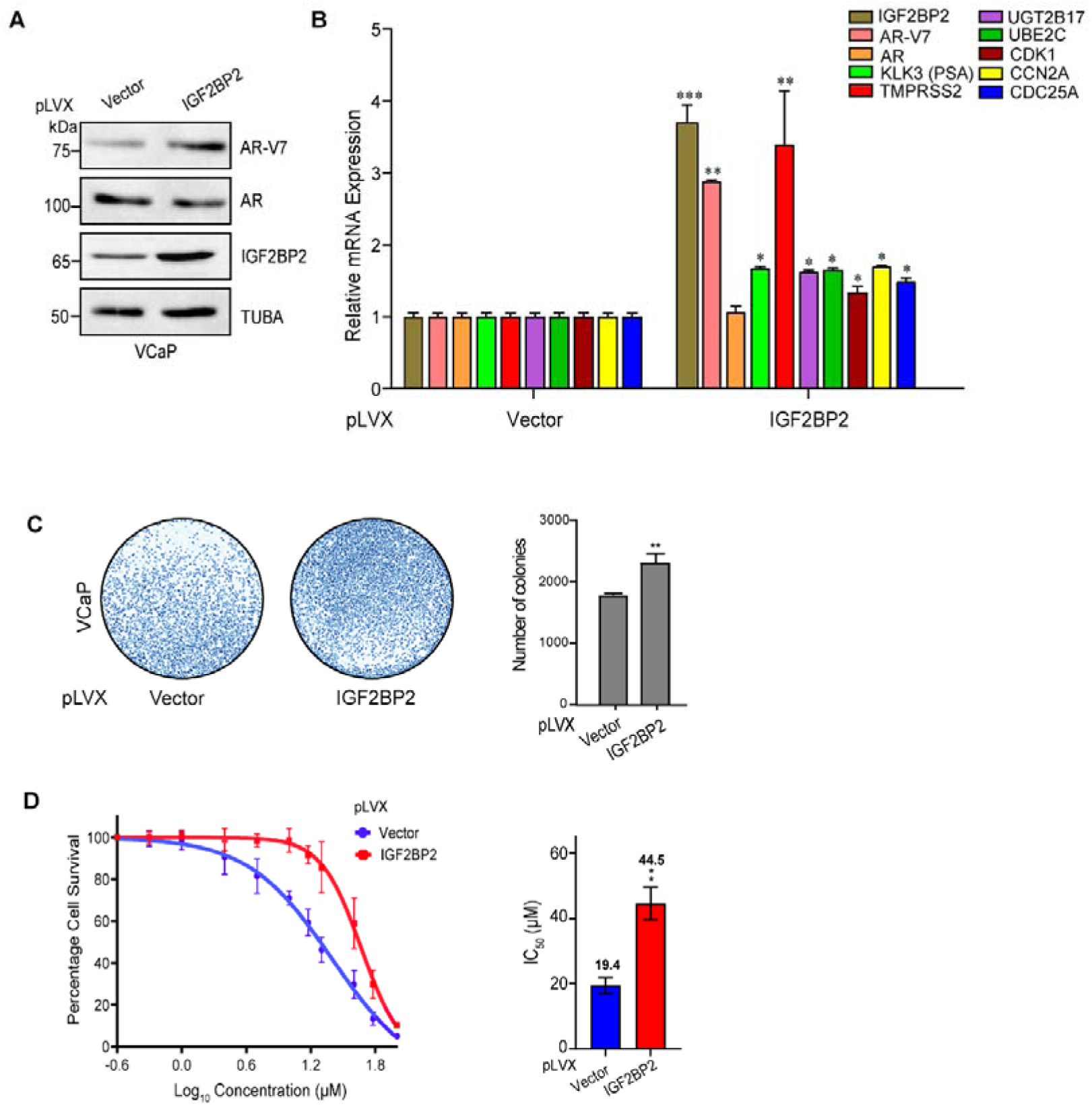
IGF2BP2 overexpression enhances AR-V7 expression and enzalutamide resistance. **A.** Immunoblot analysis of indicated proteins in IGF2BP2-overexpressing VCaP cells relative to vector-overexpressing cells. Tubulin was used as loading control **B.** qRT-PCR analysis showing mRNA expression of indicated genes in IGF2BP2-overexpressing VCaP cells relative to vector-overexpressing cells. **C.** Colony formation assay in IGF2BP2-overexpressing VCaP cells relative to vector-overexpressing cells. Colonies were stained using crystal violet, counted (n=3 biological replicates/group) and mean ± SD (\*\**p* < 0.01) was plotted. **D.** MTT assay to determine IC50 of enzalutamide in IGF2BP2-overexpressing VCaP cells relative to vector-overexpressing cells. The data is shown as the meanL±LSD; *n* = 3 independent experiments, two-tailed Student’s t-test, \*\**p* < 0.01.

These findings underscore the role of IGF2BP2 in modulating AR-V7 expression, influencing downstream target genes, and contributing to enzalutamide resistance in CRPC cells.

### 2.5 KH3 and KH4 domains of IGF2BP2 are critical for AR-V7 expression

IGF2BP2 protein comprises of two RNA recognition motifs (RRM) and four K homology (KH) domains that govern its RNA binding properties. (Fig. 5A). To pinpoint the domain(s) that might be crucial for modulating AR-V7 expression, we systematically constructed IGF2BP2 domain-deletion mutants (i.e., two RRM domains, and the four KH domains) (Fig. 5A). Immunoblot analysis in VCaP cells expressing the various deletion constructs revealed that the cells expressing IGF2BP2 protein lacking KH3 and KH4 domains fail to upregulate AR-V7 expression relative to the WT, KH1, KH2, RRM1, and RRM2 deletion expressing cells (Fig. 5B). Importantly, expression of these mutants had no discernible effect on AR expression, pointing towards the specificity of KH3 and KH4 domains in modulating AR-V7 expression.

**Figure 5:**
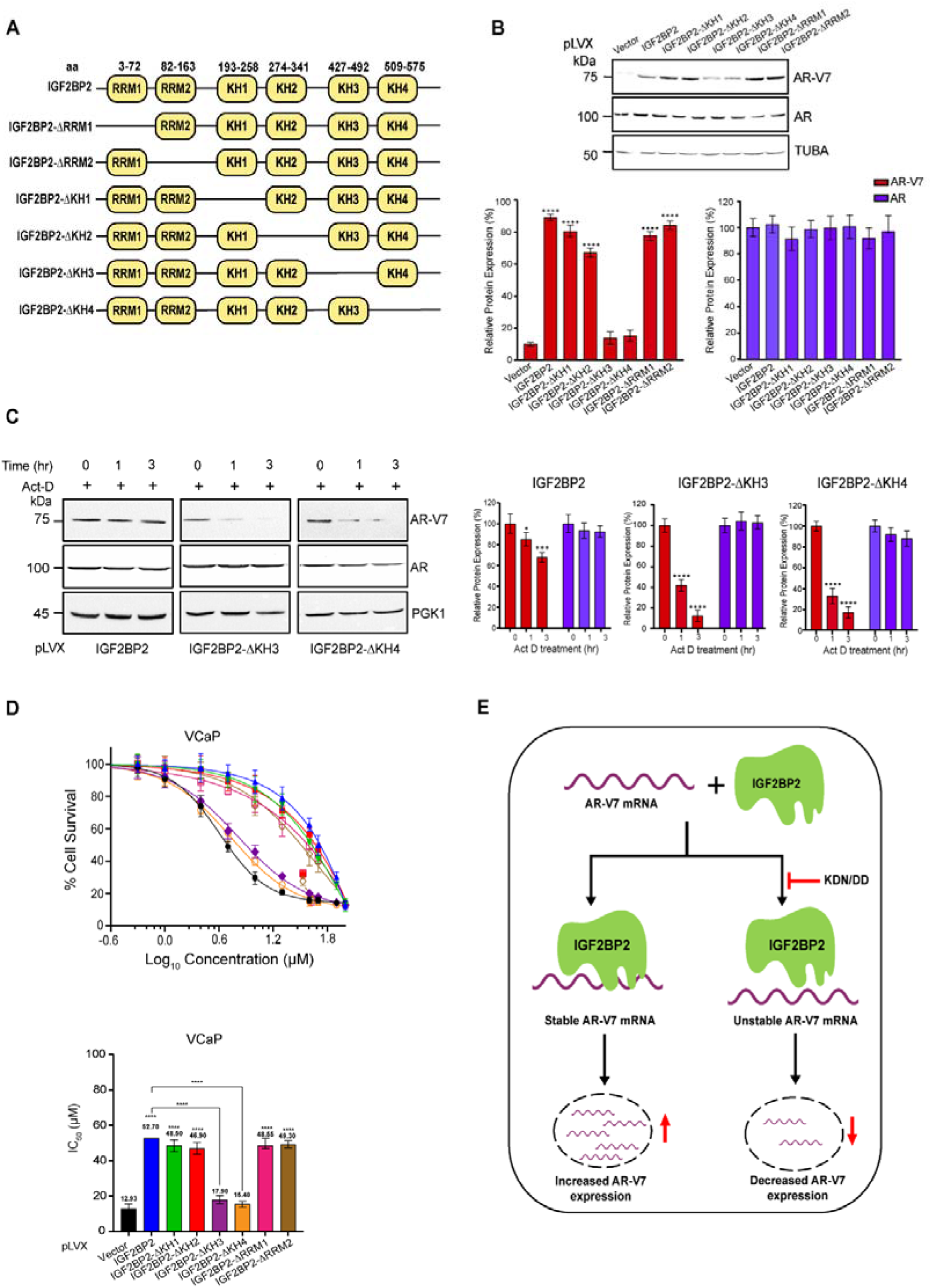
KH3 and KH4 domains of IGF2BP2 are critical for AR-V7 expression. **A.** Schematic representation of different domains present in IGF2BP2. **B.** Immunoblot analysis of AR-V7 and AR in VCaP cells overexpressing indicated domain deleted IGF2BP2 proteins relative to vector-overexpressing cells. Tubulin was used as loading control. Densitometric analysis for relative AR-V7 (red bars) and AR (blue bars) expression was performed using Image J software and normalised to TUBA expression. Data are the mean± SD. \**p* < 0.05, \*\**p* < 0.01, *** *p* < 0.001. **C.** Immunoblot analysis of indicated proteins in VCaP cells expressing either IGF2BP2-KH3 or -KH4 domain deletion proteins relative to IGF2BP2-expressing cells following actinomycin-D (5μg/mL) treatment for 0, 1 and 3hr. (n=3 biological replicates/group). Densitometric analysis for relative AR-V7 (red bars) and AR (blue bars) expression was performed using Image J software and normalised to PGK1 expression. Data are the mean± SD. \**p* < 0.05, \*\**p* < 0.01, *** *p* < 0.001. **D.** MTT assay to determine IC50 of enzalutamide in VCaP cells expressing indicated domain deleted IGF2BP2 proteins relative to vector-expressing cells. DMSO was used as control. The data are shown as the meanL±LSD; *n* =L3 independent experiments, and two-tailed Student *t* test was used to evaluate statistical the significance. \**p* < 0.05, \*\**p* < 0.01, *** *p* < 0.001, **** *p* < 0.0001 **E.** Model summarizing the effect of IGF2BP2 on AR-V7 mRNA stability.

Next, we wanted to check if KH3 and KH4 domains are also responsible for maintaining AR-V7 mRNA stability. We treated VCaP cells expressing various domain deletion constructs with actinomycin D for 0, 1 and 3hr and performed immunoblot analysis. The results revealed ∼50-60% decrease in AR-V7 protein expression 3 hours post actinomycin D treatment in cells expressing KH3 and KH4 deletion constructs relative to WT, KH1, KH2, RRM1, and RRM2 deletion expressing cells (Fig. 5C and Fig. S5). Consistent with earlier observations, none of the deletions had any impact on AR stability (Fig. 5C and Fig. S5). These results suggest that IGF2BP2 binds to AR-V7 mRNA through its KH3 and KH4 domains to regulate its stability and expression.

Next, we investigated if presence of KH3 and KH4 domains in IGF2BP2 plays a role in imparting enzalutamide resistance to CRPC cells. For this, we treated VCaP cells expressing vector, WT-IGF2BP2 and various domain deletion constructs with varying concentrations of enzalutamide and measured the IC50 using MTT assay. Notably, there was a marked increase in IC50 value for enzalutamide in cells expressing IGF2BP2, RRM1, RRM2, KH1 or KH2 deletion constructs while the cells expressing KH3 or KH4 domain deletion mutants did not exhibit any significant increase in IC50 values relative to vector-expressing cells. The IC50 value for enzalutamide IGF2BP2-expressing cells was 52.7μM, however it decreased to 17.9μM and 15.4μM in KH3 and KH4 overexpressing cells which was comparable to 12.93μM in vector-expressing cells (Fig. 5D). Collectively, these results demonstrate that the presence of KH3 and KH4 domains in IGF2BP2 is critical for regulating AR-V7 stability, expression, and enzalutamide resistance in CRPC cells.

## 3. Discussion

The aggressiveness of prostate cancer, leading to metastatic lesions and CRPC development, underscores the necessity for understanding the signaling pathways and molecules driving progression and drug resistance. Previous studies have highlighted the role of LBD-truncated AR-Vs in CRPC (7,31), with the constitutively active splice variant AR-V7 being a major driver of tumorigenesis and resistance to anti-androgens like enzalutamide and abiraterone highlighting the urgency to target AR-V7 (3,31).

IGF2BP2 belongs to the family of IGF2 RNA binding proteins, which are critical regulators of growth and development in various cancer types. Recent studies have demonstrated its role as an m6A reader, playing a causal role in the development and progression of different cancers including hepatocellular carcinoma, glioblastoma, liposarcoma, pancreatic carcinoma and ovarian carcinoma (20–22). It comprises of six highly conserved RNA binding domains (RBDs) that play an important role in the recognition, binding and stabilisation of IGF2BP2-RNA complexes and also mediate the recruitment of other RBPs (24). However, the role of IGF2BP2 in CRPC development and progression has not been previously reported. Our TCGA data analysis demonstrated that IGF2BP2 is significantly upregulated in CRPC patient samples suggesting its association with disease progression and poor survival. Based on the previous studies identifying the role of IGF2BP2 in various cancer types and our analysis of both primary prostate and CRPC patient samples, we postulated that the upregulation of IGF2BP2 might be linked to CRPC progression and enzalutamide resistance.

Here, we have discovered that IGF2BP2 downregulation significantly reduces AR-V7 inside cells without affecting the AR levels, consequently affecting CRPC progression. Moreover, the expression of AR-V7 target genes is also abrogated upon IGF2BP2 silencing suggesting the broader impact of IGF2BP2 on CRPC progression. Furthermore, we show that IGF2BP2 directly and selectively binds to the ISE region of AR-V7 mRNA and regulates its stability which is line with previous reports showing direct binding of IGF2BP2 to RNA (32,33). The enhanced mRNA stability leads to persistent AR-V7 expression inside cells, thereby contributing to CRPC progression and enzalutamide resistance, in line with previous studies demonstrating the role of IGF2BP2 in regulating cancer progression by stabilizing various mRNA and lncRNA transcripts (32–34).

Interestingly, our results also demonstrate that IGF2BP2-mediated regulation of AR-V7 expression and stability depends upon the presence of intact KH3 and KH4 domains in IGF2BP2. This points towards the ability of these domains to directly bind to AR-V7 mRNA, which is consistent with the ability of KH domains to confer sequence-specific RNA recognition ability to IGF2BP2 and reports showing that IGF2BP1 regulates mRNA stability of a large number of targets via its KH3KH4 domains (24,35). Interestingly, we observe that the presence of intact KH3 and KH4 domains also positively correlates with enzalutamide resistance, highlighting the importance of sequence-specific interaction of these domain scaffolds with AR-V7 mRNA in regulating drug resistance and disease progression. This observation is also consistent with previous studies demonstrating that IGF2BP1 regulates m6A methylation of target transcripts in cancer cells via KH3KH4 di-domain (24).

In summary, our results, for the first time, demonstrate that IGF2BP2 regulates AR-V7 expression by directly binding to the AR-V7 pre-mRNA. Altering IGF2BP2 expression prevents tumor cell proliferation and resensitizes the cells to androgen deprivation therapy. Our mechanistic analysis reveals that KH3 and KH4 domains in IGF2BP2 are responsible for its interaction with AR-V7 mRNA thereby helping regulate its stability. Taken together, the identification of IGF2BP2 as a key modulator of AR-V7 expression positions it as a potent therapeutic target for future drug discovery efforts.

## Limitations and future directions

Our study sheds light on the role of IGF2BP2 in regulating AR-V7 stability and its implication in enzalutamide resistance. However, additional studies would be required to precisely elucidate the specific sequence(s) within the AR-V7 ISE region that are bound by IGF2BP2 to regulate its stability. Future experiments involving comprehensive mutagenesis of AR-V7 mRNA will be required to pinpoint the exact binding sequence(s) of IGF2BP2 and its KH3 and KH4 domains on AR-V7 mRNA. Elucidating IGF2BP2 crystal structure in complex with AR-V7 mRNA could further aid these experiments and provide detailed insights into the mechanism of action of IGF2BP2. Moreover, while our cell line studies demonstrate that IGF2BP2 inhibition can mitigate oncogenic AR-V7 signaling and enzalutamide resistance, it would be worth corroborating these findings in *in vivo* models of CRPC.

## 4. Materials and Methods

### Cell culture

HEK293T, VCaP, and 22Rv1 cells were purchased from ATCC. LnCaP95 cells were a kind gift from Dr. Jun Luo, Johns Hopkins School of Medicine, Baltimore, MD. HEK293T and VCaP cells were cultured in DMEM media (HiMedia), and 22Rv1 cells were cultured in RPMI1640 media (HiMedia) supplemented with Pen-strep antibiotics, L-glutamine and 10% FBS . LnCaP95 cells were cultured in Phenol-red free RPMI1640 media (Himedia) supplemented with Pen-strep antibiotics, L-glutamine and 10% charcoal-stripped FBS (CFBS). All the cell lines were maintained in presence of 5% CO2 at 37°C and were tested for any bacteria, fungi and mycoplasma contamination.

### Viral Production and Generation of stable knockdown cells

For lentivirus production, HEK293T cells were seeded in individual wells of a 6-well plate and incubated for 24h at 37°C after which they were co-transfected with lentiviral packaging plasmids along with IGF2BP2 shRNA plasmid using Effectene as a transfection reagent. 48h post-transfection, the viral supernatant was filtered using a 0.45μM filter. For stable knockdown of IGF2BP2, cells were infected either with non-silencing shRNA or IGF2BP2 shRNA. Forty-eight hours post-infection, cells were selected using appropriate puromycin concentrations (0.5μg/mL for 22Rv1 and 2μg/mL for VCaP and LnCaP95) for 4-5 days. IGF2BP2 shRNA sequences used in this study are listed in Supplementary Table 1.

### RNA preparation, cDNA preparation and quantitative PCR (qPCR) analysis

Total RNA was isolated from cells using TRIzol (Invitrogen, USA) and cDNA synthesis was performed using the Superscript II cDNA Synthesis Kit (Invitrogen, USA). qRT-PCR was performed using the PowerUp SYBR Green Master Mix, with gene-specific primers as per manufacturer’s instructions. Expression of target mRNAs was quantified using ΔΔCT method relative to the house-keeping gene ACTB. Primer sequences for all the genes analysed in this study are listed in Supplementary Table 2.

### Protein extraction and Immunoblot analysis

For total protein cell lysate preparation, cells were harvested using cell scraper and lysed for 10 minutes using RIPA buffer (20mM Tris-HCl (Affymetrix), pH-7.5, 150mM NaCl (Affymetrix), 1mM EDTA (Affymetrix), 1mM EGTA (HiMedia), 1% NP-40, 1% Sodium deoxycholate (Sigma), 1% sodium orthovandate (sigma) containing protease (Roche) and phosphatase inhibitor cocktail), following which it was centrifuged at 14,000 rpm for 30 min. Protein concentration in the clarified supernatant was quantified using Bradford Protein Assay Reagent (Bio-Rad Laboratories). 40μg of protein lysate was loaded onto SDS-PAGE gel using a Mini-Protean System (BioRad) and transferred onto nitrocellulose membrane (BioRad 0.2μm) for 3h at 250mA. The membranes were blocked in Tris buffered saline containing 0.1% Tween 20 (TBST) containing 5% skimmed milk for 1h following which primary antibodies (AR-V7: AG10008 (Precision antibody); AR: 06-680 (Merck Millipore) ; PGK1: ab38007 (abcam) Tubulin: ab15246 (abcam) ) were added overnight at 4°C. After probing with primary antibody, membranes were washed using TBST and then incubated with HRP-conjugated secondary antibody at room temperature for 1h. Blots were finally developed using chemisluminescent substrate (SuperSignal West Pico PLUS Chemiluminescent Substrate, Thermo Fisher Scientific) in chemiDoc imaging system (Bio-Rad Laboratories).

### Plasmid construction

Human IGF2BP2 was cloned into p3XFLAG-myc-CMV 26 (Sigma) and pLVX-E1Fα-IRES-mCherry-Puro (Takara Bio) to perform the experiments associated with its ectopic expression. RRM domain truncations (IGF2BP2-ΔRRM) and KH domain truncations (IGF2BP2-ΔKH) were generated by performing overlap PCR reaction (1) using primers listed in supplementary table 3 and cloned in p3XFLAG-myc-CMV 26 and pLVX-E1Fα-IRES-mCherry-Puro between EcoRI and XbaI sites. All the PCR products and clones were verified by DNA sequencing.

### mRNA stability assay

Cells infected with either NS, IGF2BP2 shRNA or various IGF2BP2-expression plasmids were seeded in individual wells of a 6-well plate and incubated till 80% confluency was achieved. Subsequently, the cells were treated with 5μg/mL actinomycin D (sigma) and harvested at 0h, 1h and 3h timepoints. The whole cell protein lysate was prepared and analysed using western blotting. The protein degradation was quantified by calculating the band intensity using densitometry analysis. All the data was derived from three independent biological replicates.

### RNA Immunoprecipitation and qRT-PCR analysis

IGF2BP2 overexpressing VCaP cells were washed with ice-cold PBS post-infection followed by lysis of cells using immunoprecipitation (IP) lysis buffer containing 50 mM Tris-Cl (pH 7.4), 250 mM NaCl, 5 mM EDTA, 0.5 mM DTT, 0.5% Triton X-100, 1X protease inhibitor (Roche) ) on ice, following which the lysate was subsequently centrifuged at 14,000 x g for 30 min at 4°C. The clarified supernatant was carefully removed and 1mg/sample lysate was incubated with 1ug of indicated antibodies at 4°C. IgG antibody was used as negative control. 20μL Protein G-coated Dynabeads (Life Technologies) were added to the mixture, incubated at 4°C for 2h, separated using magnetic stand and washed 3X times with IP lysis buffer. Bound RNA was eluted from the dynabeads using TRIzol, reverse transcribed and the enrichment of IGF2BP2 on ESE and ISE regions was estimated using qRT-PCR. The primers used for the analysis are listed Supplementary Table 2.

### Wound Healing Assay

VCaP cells were infected with indicated shRNAs in a 6-well plate and incubated until 80% confluency was achieved. After the cells formed a confluent monolayer, scratches were performed using 200μL tip and detached cells were removed by giving PBS wash following which fresh culture media was added. The closure of scratch was analysed under the microscope until the scratch was completely covered in the control well. Images were taken under the microscope (Nikon DS-F*i*3 microscope). Difference in wound width was measured to calculate the cell migration ability. All the data was derived from three independent biological replicates.

### Colony formation assay

5X10^3^, VCaP, LnCaP95 or 22Rv1 cells infected with various viruses were seeded in individual wells of a 6-well plate and incubated at 37°C until distinct colonies were formed. Cells were washed with PBS and stained with 0.01% crystal violet. Positively stained colonies were observed under the microscope and imaged.

### Soft agar assay

The effect of IGF2BP2 knockdown on anchorage-independent growth of 22Rv1 and LnCaP95 cells was monitored in a 6-well plate coated with basal layer of 1.2% Nobel agar and RPMI-1640 containing 20% FBS. Stable knockdown cells were seeded at 5000 cells/well density in media mixed with 0.6% Nobel agar. After the agar was set, 1mL RPMI 1640 + 10% FBS media was added to each well. 3 weeks later, the macroscopic colonies formed on the plate were stained with 0.001% crystal violet and imaged using a phase contrast microscope with a 10X objective.

### Cell proliferation assay

5 X 10^3^, 22Rv1 or LnCaP95 cells (NS and IGF2BP2 knockdown) were seeded in each well of a 96-well plate with a density. Following 24h incubation, cells were treated with varying concentrations of Enzalutamide (0.5μM - 100μM) using DMSO as a control. Cell proliferation post-enzalutamide treatment in IGF2BP2 knockdown cells was analysed using MTT reagent, and the absorbance value was measured at OD 595nm using a microplate reader (BioRad). Each group was analysed in triplicate.

### Statistical Analysis

Statistical analysis was performed using t-test (two-tailed unpaired) and ANOVA in GraphPad Prism (www.graphpad.com). All the experiments included at least three biological replicates unless stated otherwise. Data is represented as mean ± S.D. and the statistical significance is denoted by *p*-values.

## Supplementary material

Supporting information and legends are available separately.

## Author contributions

Conceptualization, Taruna Saini, Parul Mishra and Ashish Misra; Formal analysis, Taruna Saini, Devesh Srivastava and Rajnikant Raut; Funding acquisition, Ashish Misra; Investigation, Taruna Saini; Methodology, Taruna Saini; Project administration, Parul Mishra and Ashish Misra; Resources, Ashish Misra; Supervision, Parul Mishra and Ashish Misra; Writing – original draft, Taruna Saini, Rajnikant Raut, Parul Mishra and Ashish Misra; Writing – review & editing, Taruna Saini, Devesh Srivastava, Parul Mishra and Ashish Misra.

## Acknowledgments

This work is supported by ICMR (2021-13073) to A.M., HGK-IYBA (BT/11/IYBA/2018/08) and UoH-IoE-RC5-22-011 grants to P.M.. T.S. is a recipient of MoE-GoI fellowship. R.R. duly acknowledges CSIR-SRF fellowship support. D.S. is a recipient of PMRF. The authors sincerely thank Dr. Jun Luo, Johns Hopkins School of Medicine for sharing LnCaP95 cells for our studies.

## Institutional Review Board Statement

Not applicable.

## Informed Consent Statement

Not applicable.

## Data Availability Statement

All data generated during this study are included in this published article and its supplementary files.

