## Supplementary material for "Insulin-like Growth Factor 2 mRNA-binding protein 2 (IGF2BP2) Promotes Castration-Resistant Prostate Cancer Progression by Regulating AR-V7 mRNA Stability": Fig. S1, Fig. S2, Fig. S3, Fig. S4, Fig. S5

**Supplementary Figures**

**Figure S1**

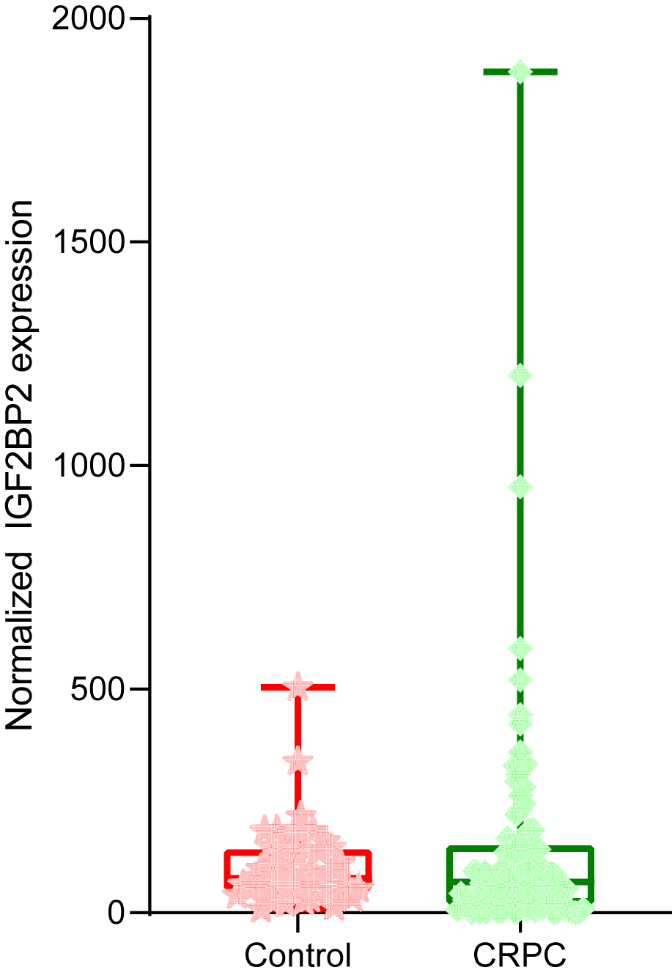

Figure S1. Comparative expression of IGF2BP2 in normal patients (n= 52) versus metastatic CRPC patients (n=99) from TCGA dataset (*p*=0.031 and logFC=0.7). Gene expression is graphically represented as box and whiskers plot with outliers.

**Figure S2**

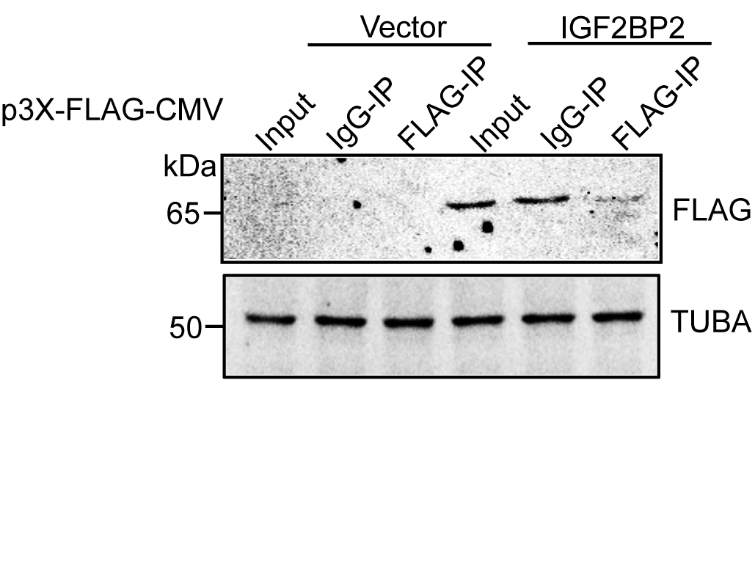

Figure S2. Western blot analysis in the flow-through after anti-FLAG immunoprecipitation in VCaP cells were transiently transfected with either FLAG-IGF2BP2 or empty vector for 72 hours. All immunoprecipitation experiments were performed in biological triplicates.

**Figure S3**

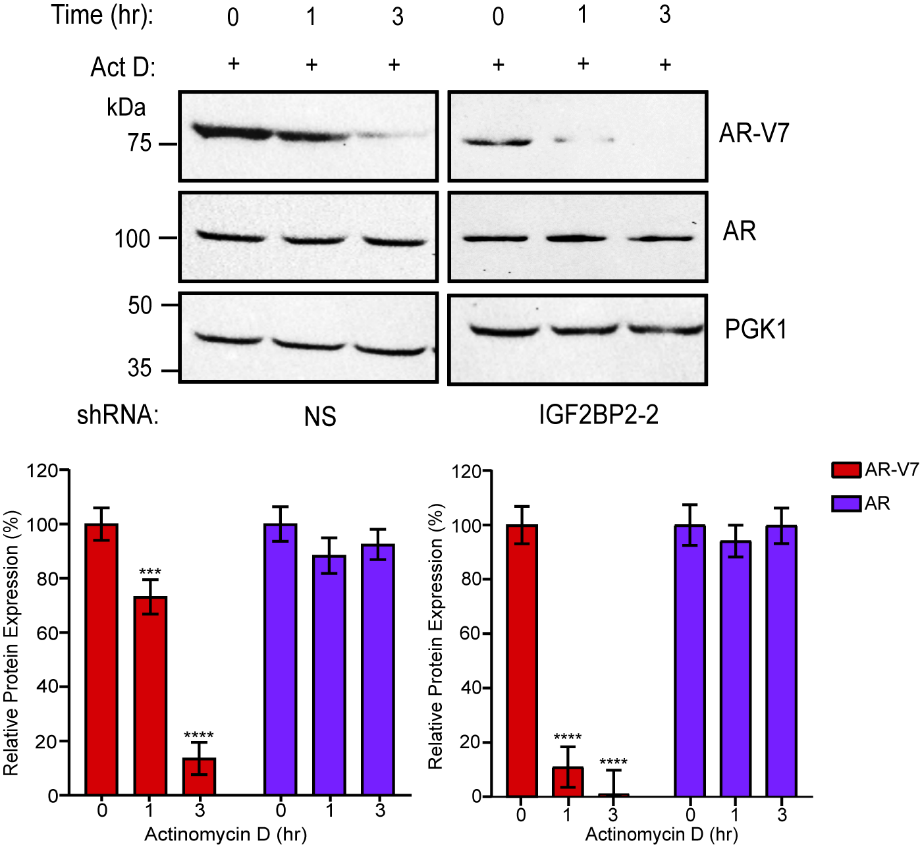

Figure S3. Expression of AR-V7, AR and PGK1 was determined by immunoblot analysis after Actinomycin D treatment (5 μg/mL) at indicated time in IGF2BP2 knockdown 22Rv1 cells. Densitometric analysis of relative expression of AR-V7 and AR was performed using Image J software and normalised by the expression of control. Data are the mean± SEM. **p* < 0.05, *** p < 0.001.

**Figure S4**

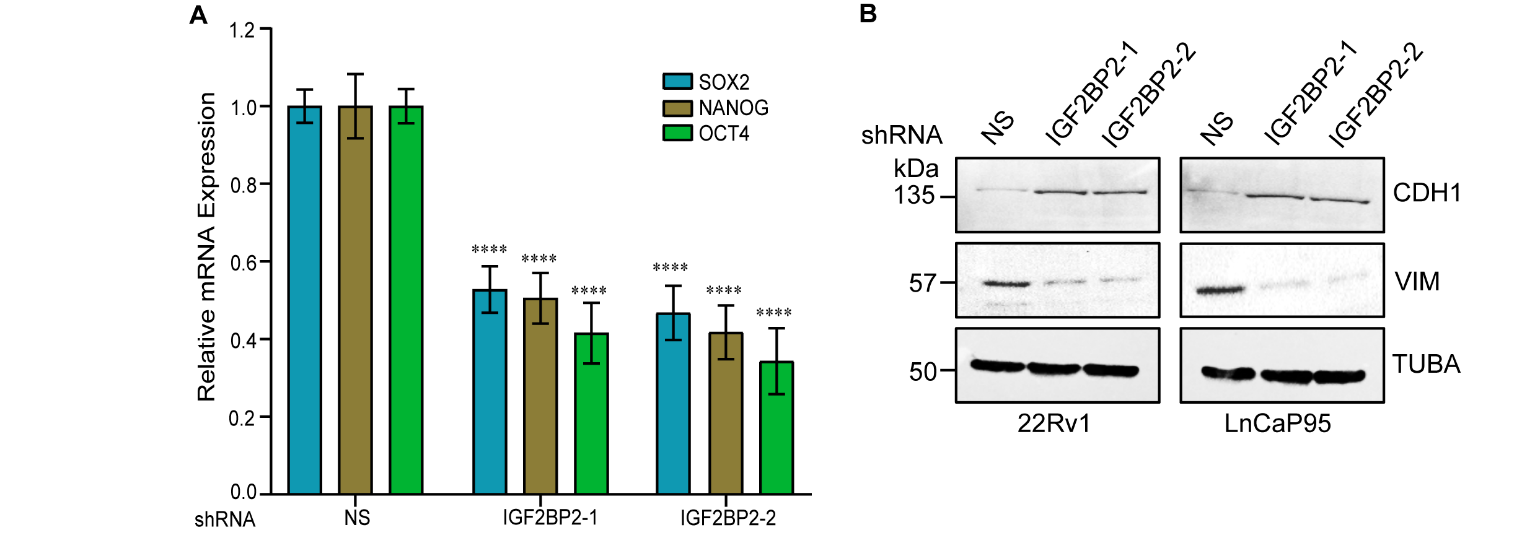

Figure S4. (A) Quantitative gene expression analysis of stem cell genes Sox2, Nanog and Oct-4 by performing qRT-PCR in IGF2BP2 depleted 22Rv1 cells; (B) Protein expression analysis of common EMT markers E-cadherin and Vimentin in IGF2BP2 knockdown 22Rv1 and LnCaP95 cells. Tubulin was used as a loading control.

**Figure S5**

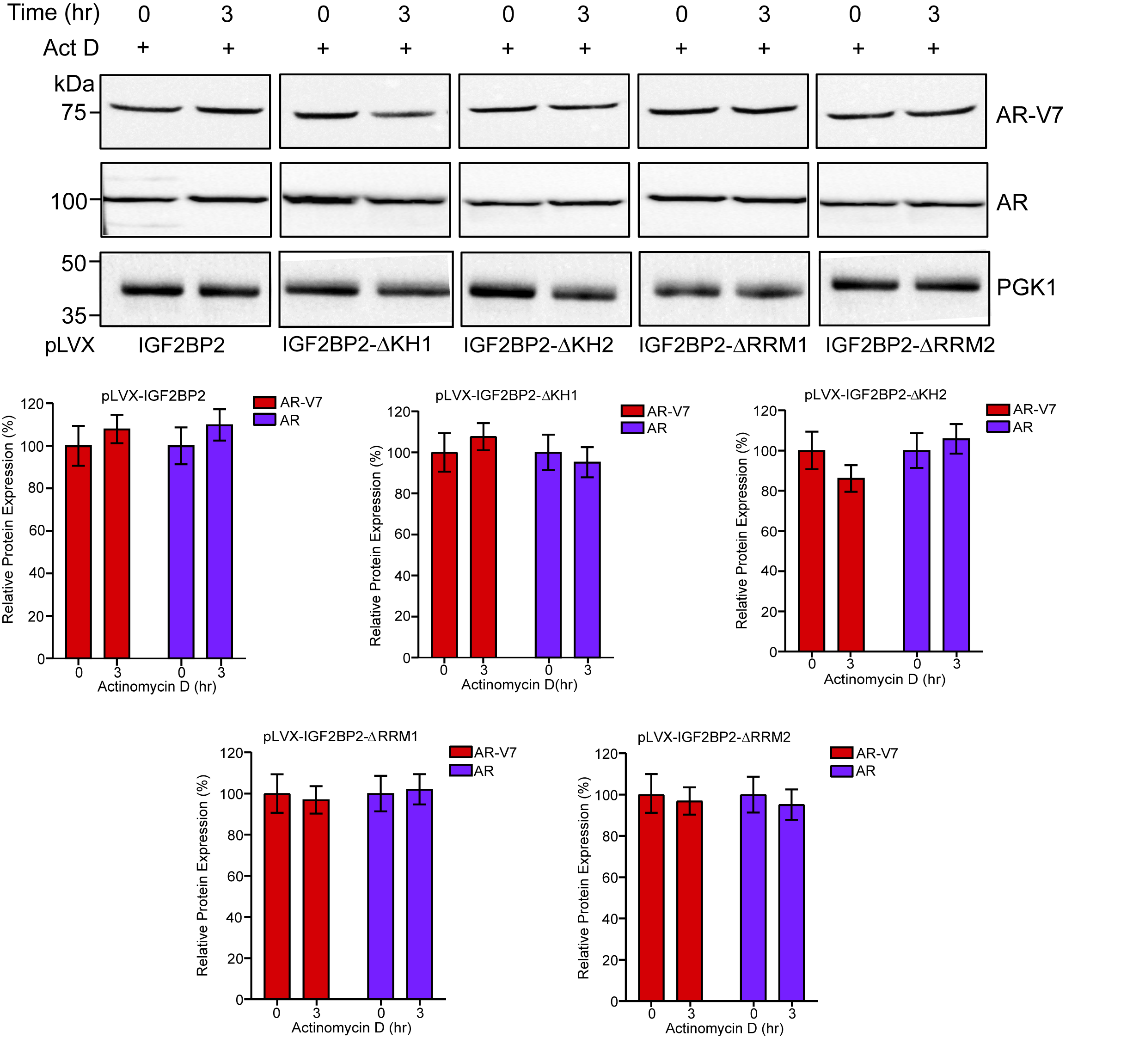

Figure S5. Consistent AR-V7 and AR protein expression on overexpressing full-length IGF2BP2 and KH1-2 and RRM1-2 truncated IGF2BP2 constructs in VCaP cells. Densitometric analysis of relative expression of AR-V7 and AR was performed using Image J software and normalised by the expression of control. Each value in the data represents mean± SEM.
