## Supplementary Table 1, Supplementary Table 2, Supplementary Table 3 for "Insulin-like Growth Factor 2 mRNA-binding protein 2 (IGF2BP2) Promotes Castration-Resistant Prostate Cancer Progression by Regulating AR-V7 mRNA Stability"

**Supplementary Tables**

**Supplementary Table 1. List of IGF2BP2 shRNA**

| **SEQUENCE** | **TARGET REGION** |
| --- | --- |
| CGGATCTTTGGGAAACTGAAA | KH3 DOMAIN |
| CTTAACCAGTGCAGAAGTCAT | KH4 DOMAIN |

**Supplementary Table 2. List of RT-qPCR primers**

| **GENE NAME** | **SEQUENCE** |
| --- | --- |
| AR | F: 5′ TCTTGTCGTCTTCGGAAATGT 3′  R: 5′ AAGCCTCTCCTTCCTCCTGTA 3′ |
| AR-V7 | F: 5′ CGGAAATGTTATGAAGCAGGGATGA 3′  R: 5′ CTGGTCATTTTGAGATGCTTGCAAT 3′ |
| IGF2BP2 | F: 5′ AGTGGAATTGCATGGGAAAATCA 3′  R: 5′ CAACGGCGGTTTCTGTGTC 3′ |
| KLK3 | F: 5′ AGTGCGAGAAGCATTCCCAAC 3′  R: 5′ CCAGCAAGATCACGCTTTTGTT 3′ |
| TMPRSS2 | F: 5′ ATGAAAACCATGGATACCAACCG 3′  R: 5′ GAGAATCCATCTTGAGGTGC 3′ |
| UBE2C | F: 5′ TGCCCTGTATAGATGTCAGGA 3′  R: 5′ GGGACTATCAATGTTGGGTTCT 3′ |
| CDC25A | F: 5′ CAAACCTTGACAACCGATG 3′  R: 5′ ACACTGACCAGTCGTGGAG 3′ |
| UGT2B17 | F: 5′ ACCCAGCCAAACCCTTGCCTAA 3′  R: 5′ GGCTGATGCAATCATGTTGGCAC 3′ |
| ELK1 | F: 5′ CAGCCAGAGGTGTCTGTTACC 3′  R: 5′ GAGCGCATGTACTCGTTCC 3′ |
| CDK1 | F: 5′ CCTAGCATCCCATGTCAAAAACTTGG 3′  R: 5′ TGATTCAGTGCCATTTTGCCAGA 3′ |
| CCN2A | F: 5′ GAAGACGAGACGGGTTGCA 3′  R: 5′ AGGAGGAACGGTGACATGCT 3′ |
| SOX2 | F: 5′ GAGCTTTGCAGGAAGTTTGC 3′  R: 5′ GCAAGAAGCCTCTCCTTGAA 3′ |
| NANOG | F: 5′ ACCTTGGCTGCCGTCTCTGG 3′  R: 5′ AGCAAAGCCTCCCAATCCCAAACA 3′ |
| OCT-4 | F: 5′ TTTTGGTACCCCAGGCTATG 3′  R: 5′ GCAGGCACCTCAGTTTGAAT 3′ |

**Supplementary Table 3. List of Cloning primers**

| **GENE NAME** | **SEQUENCE** |
| --- | --- |
| IGF2BP2 | F: 5′ CAAAAGCTTATGGCGACGGAGCATCCC 3′  R: 5′ CAAGCGGCCGCAAACTTGAACTCCTTATATTTCTTG 3′ |
| IGF2BP2-ΔRRM1 | F: 5′ CGGGAATTCATGAAAAAGCTAAGGAGCAGG 3′  R: 5′ CAAGCGGCCGCAAACTTGAACTCCTTATATTTCTTG 3′ |
| IGF2BP2- ΔRRM2 | F: 5′ AAAAAGCTAAGGAGCGAAGAGGTGAGCTCCCC 3′  R: 5′ GGGGAGCTCACCTCTTCGCTCCTTAGCTTTTT 3′ |
| IGF2BP2- ΔKH1 | F: 5′ AGGCCAGACAGATTCTTGAAATCATGCAGAA 3′  R: 5′ TTCTGCATGATTTCAAGAATCGTCTGGCCCT 3′ |
| IGF2BP2- ΔKH2 | F: 5′ GACCAAACTAGCCGAAATGAAGAAGCTGC 3′  R: 5′ GCAGCTTCTTCATTTCGGCTAGTTTGGTC 3′ |
| IGF2BP2- ΔKH3 | F: 5′ CACTCTTATCCAGAGTTTGGGAAACTGAAAGAGG 3′  R: 5′ CCTCTTTCAGTTTCCCAAACTCTGGATAAGAGTG 3′ |
| IGF2BP2- ΔKH4 | F: 5′ CCCCAAAGAAGAAGTGGAAATTGTACAACAGGTG 3′  R: 5′ CACCTGTTGTACAATTTCCACTTCTTCTTTGGGG 3′ |
